# Efficient genome graphs indexing, querying and visualization with gfaidx and gfaidx-viz

**DOI:** 10.64898/2026.11.15.766357

**Authors:** Fawaz Dabbaghie, Martin Steinegger

## Abstract

Genome graphs enable the analysis of genetic variation across many genomes, but exploring individual regions remains computationally demanding. We present gfaidx, a general-purpose indexer giving fast, low-memory access to pangenome, assembly, and de Bruijn graphs in GFA format. It retrieves gene-sized subgraphs and their haplotypes over 100-fold faster than existing tools such as ODGI and vg, and can index human pangenome-scale graphs on a personal computer. The accompanying gfaidx-viz interactively visualizes pre-indexed human pangenomes. Both are freely available at https://github.com/fawaz-dabbaghieh/gfaidx and https://github.com/fawaz-dabbaghieh/gfaidx-viz.

---

Graphs are widely used in biology to represent relationships, from interactions between proteins to connections between DNA sequences. Genome graphs are non-linear representations of DNA segments as connected nodes, which encode genetic variation as alternative paths, enabling multiple genomes to be compared within a single representation and supporting tasks from genome assembly to variant calling and pangenomics (1–4). Genome graphs are growing rapidly, for example, the Human Pangenome Reference Consortium (HPRC) expanded its pangenome graph from 94 haplotypes in its first version (5) to 464 in its second version (6).

However, many biological questions still concern a single gene, structural variant, or region of interest rather than the entire graph. Investigating such a region requires access to its sequences, connections, and the haplotypes traversing it, so ideally a query would retrieve only a small part of the graph. Yet with existing tools this often requires loading much larger graph into memory, so even local queries become computationally demanding. The need for such local access extends beyond human pangenomes. De Bruijn graphs, which connect overlapping sequences, support genome and metagenome assembly (1) and more recently, allow for search over petabase-scale sequence collections (7). In bacterial genome-wide association studies, for example, extracting local subgraphs around phenotype-associated sequences helps interpret their genomic context (8).

In the linear-reference world, this problem is already solved: indexed, block-compressed formats such as `faidx, tabix,` and BAM/CRAM, together with the `HTSlib` library, let applications retrieve regions without loading entire datasets and power interactive genome browsers such as IGV (9–12). For graphs, on the other hand, subgraph queries with tools such as vg (13, 14) and ODGI (15) incur the cost of loading the whole graph into memory, making local queries expensive. More recently, `GBZ-base` (16) introduced disk-backed queries by storing a GBZ graph (17) in SQLite, which avoids loading the whole graph. However, it is specific to the GBZ representation and cannot identify the other haplotypes traversing a queried region or determine their coordinates, as both GBZ and `GBZ-base` currently lack efficient data structures for that (16).

Here, we present `gfaidx,` a graph indexer that enables fast, low-memory random access to graphs stored in Graphical Fragment Assembly (GFA) format (18), including pangenome, assembly, de Bruijn and reference rGFA graphs, with or without embedded paths or walks. Both index construction and querying operate primarily on disk, so human pangenome-scale graphs can be indexed and explored on a personal computer rather than a high-memory compute node. Users can retrieve an individual node, a local neighborhood or a genomic interval along indexed walks, resolving every walk to its named haplotype and coordinates when available. We provide C++ and Python interfaces and an interactive web visualizer, `gfaidx-viz,` for exploring pre-indexed HPRC v1 and v2 graphs.

As shown schematically in Fig. 1a, `gfaidx` iteratively partitions the graph into communities using the Louvain method (19). Each community is compressed as an independently accessible gzip member. Sidecar indices map node identifiers, paths, walks and sample coordinates to byte ranges (Methods). A query can be a node, a subgraph or a genomic region over a walk or path Fig. 1b; in each case, `gfaidx` reads only the relevant index entries and decompresses only the community or communities it spans. Thus, query cost depends primarily on the queried subgraph and the number of communities it touches, rather than on the size of the full graph.

**Fig 1.**
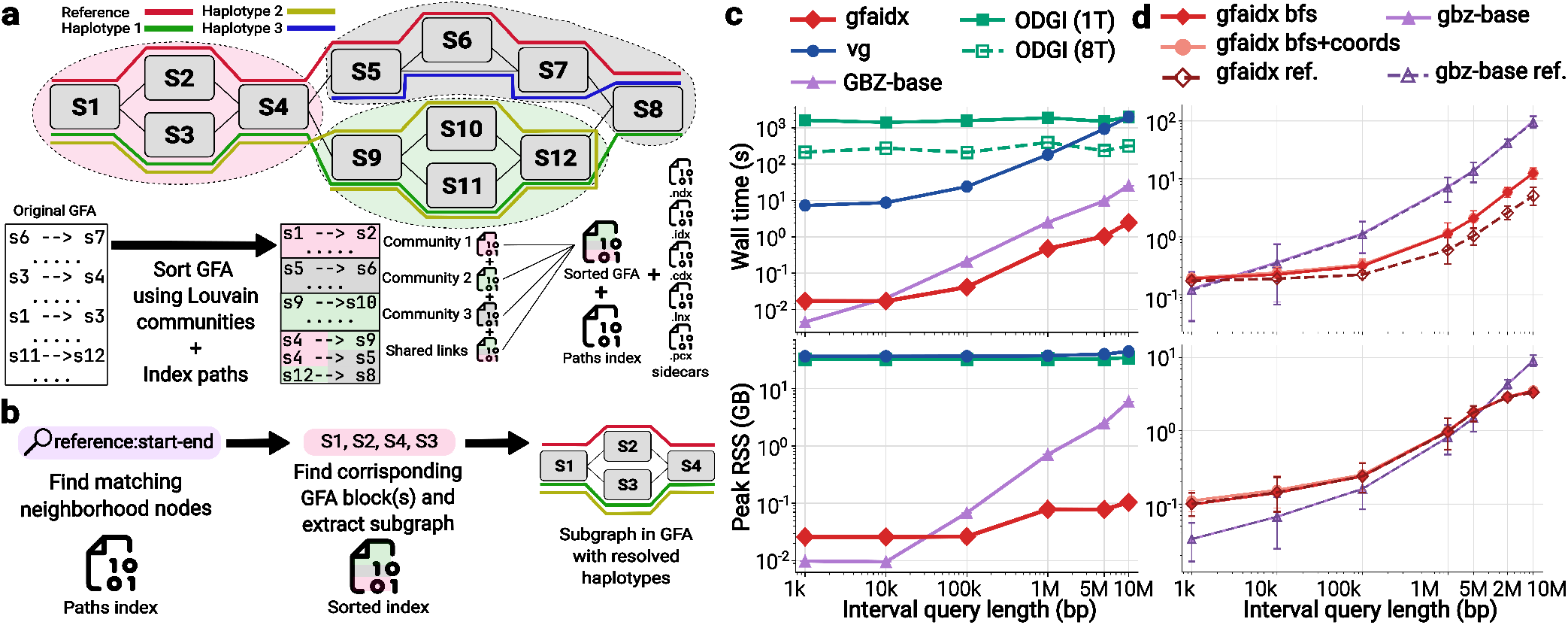
**(a)** The graph is partitioned into communities using the Louvain method, and the GFA is reordered so that each community forms an independently decompressible block. If the graph contains paths or walks, these are also indexed, and sidecar files are produced for fast random access. **(b)** Region query pipeline. The path/walk index is first used to find the nodes matching the query and to extract alternative paths; e.g., the reference query here matches S1, S2 and S4, but S3 is also retrieved. Only the block(s) of the multi-member compressed GFA containing these nodes are then accessed to produce a GFA subgraph with all haplotypes enumerated. **(c)** Wall time (top) and peak memory (bottom) for extracting the subgraph of a CHM13 interval (1 kb–10 Mb) from the HPRC v2 Minigraph-Cactus chromosome 1 graph. Settings: `gfaidx` haplotype-span extraction (`get_region –mode all –with_coords`), `ODGI extract -r` with default path merging (1 and 8 threads), `vg chunk -c 0` and `GBZ-base query –context 0`. **(d)** Subgraph extraction from the complete HPRC v2 Minigraph-Cactus graph for 1 kb–10 Mb intervals at 12 loci (Methods), with 10 kb of context (returning the surrounding subgraph and all its walks). `gfaidx` is shown with the reference walk only (ref.), all walks (bfs) and all walks with coordinates (bfs + coords); `GBZ-base` with the reference walk only and with all walks (no names or coordinates). Points are means across loci (median of runs 2–3 per locus); error bars are bootstrap 95% confidence intervals.

On the HPRC v2 Minigraph-Cactus chromosome 1 graph, `gfaidx` built its index in about 26 minutes using ~3 GB of RAM, compared with ~1.5 hours and ~345 GB for `vg’s` XG graph and 40 minutes and 83 GB for `ODGI’s` OG graph. `GBZ-base` built its database in only 32 seconds using ~1.5 GB, but it requires a GBZ graph, which took vg a further 30 minutes and ~37 GB to build. For the complete HPRC v2 Minigraph-Cactus graph, `gfaidx` built its index directly from the GFA in about 6 hours using ~40 GB on 8 threads, whereas `GBZ-base` needed 14 minutes and ~12 GB to build its database from the precomputed HPRC GBZ graph;this excludes the cost of building the GBZ graph with `vg.` To evaluate query performance, we benchmarked `gfaidx` against `vg, ODGI,` and `GBZ-base` on the HPRC v2 Minigraph-Cactus chromosome 1 graph, measuring the time and peak memory for extracting subgraphs over reference intervals of increasing size (Fig. 1c). Here, `gfaidx` returned subgraphs over a genomic interval in well under a second for smaller intervals using less than 150 MB of memory, whereas ODGI needed on the order of 10^3^ seconds per query and `vg` several seconds to 10^3^ seconds, both using more than 40 GB of memory. The disk-backed `GBZ-base` was faster for the 1 kb interval but about 10 times slower at 10 Mb, and its memory use grew with the interval size (Fig. 1c, bottom).

We next benchmarked the two disk-backed tools, `gfaidx` and `GBZ-base,` on the complete HPRC v2 Minigraph-Cactus graph (233 samples, 464 haplotypes), extracting 1 kb to 10 Mb intervals from 12 complex loci (Fig. 1d, Methods). We introduced a reference-only mode to `gfaidx` and a context BFS mode to match `GBZ-base,` because, by default, `gfaidx` calculates all the haplotypes with their coordinates for a region query. Both tools returned identical subgraphs. `GBZ-base` was faster for intervals up to 1 kb, above which `gfaidx` was faster. At 10 Mb, `gfaidx` returned the complete subgraph in approximately 12.5 seconds versus 99 seconds for `GBZ-base` (around 5 versus 95 seconds in reference-only mode), using less memory. `GBZ-base` returns the subgraph’s walks unresolved, i.e., anonymous, without haplotype names or coordinates, whereas `gfaidx` resolves every walk to its named haplotype with sub-coordinates, and does so at negligible additional cost.

To allow users to utilize the index programmatically in their own software, `gfaidx` also offers a C++ library and a Python package `(pygfaidx)`. To demonstrate the benefit of the C++ library, we integrated it in `BandageNG` (20, 21), enabling Ban-dageNG to query the indexed graph and extract subgraphs for visualization natively in seconds, instead of loading and visualizing the complete graph at once (Methods, Supplementary Fig. S2). Moreover, we integrated `pygfaidx` into `BubbleGun` (22), enabling superbubble detection without loading the full graph (Methods) and reducing memory use more than 10-fold on the HPRC v2 chromosome 1 Minigraph-Cactus and PGGB graphs (Supplementary Table S2). We also integrated `gfaidx` into the `Foldseek` webserver API (23), and connected it to a WebAssembly build of Bandage we call `gfaidx-viz;` the app provides the pre-indexed HPRC graphs for the users to query and visualize on demand (Methods).

We demonstrate the capabilities of `gfaidx` and `gfaidx-viz` in practice by querying three clinically important loci across all 464 haplotypes of the complete HPRC v2 Minigraph-Cactus graph and coloring the subgraph nodes by gene (Methods). `gfaidx-viz` renders both a linear and a haplotype (force-directed) view of the same subgraph on demand. We first queried the amylase locus. The linear view calculates the copy number per haplotype directly from the graph in comparison to the CHM13 reference, showing a haplotype carrying four copies of *AMY1*, the salivary-amylase gene whose copy-number rise is associated with high-starch diets (Fig. 2a) (24, 25). Coloring nodes by gene also lets `gfaidx-viz` show how a graph handles paralogs. At the Rh blood-group locus, *RHD* and *RHCE* occupy disjoint node sets, so the common *RHD* deletion that causes the RhD-negative type appears directly as haplotype walks that bypass the *RHD* loop (Fig. 2b) (26). The same coloring exposes the opposite case at the complement *C4* locus, where the near-identical paralogs *C4A* and *C4B* share 44% of their nodes, yet the HERV-K(C4) insertion is still resolved as a loop that only some haplotypes traverse (Supplementary Fig. S1) (27, 28).

**Fig 2.**
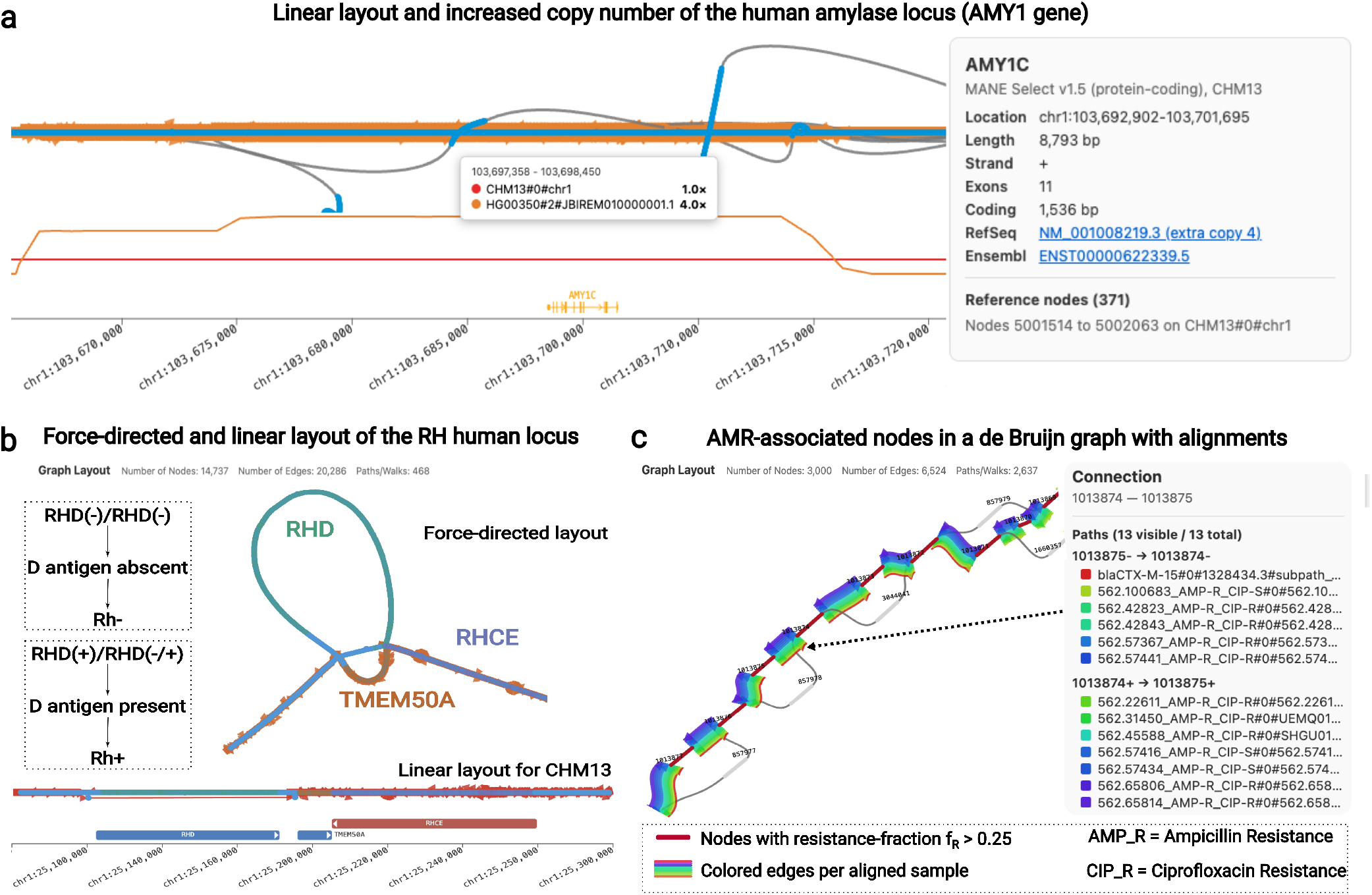
Loci extraction and visualization with gfaidx. All panels represent graphs extracted with `gfaidx` and visualized with `gfaidx-viz`. **(a)** The amylase locus (chr1:103.55–103.81 Mb) in linear layout; the copy-number track compares HG00350 haplotype 2 (orange) with CHM13 (red), showing the increase in copy-number on the *AMY1C* gene. The panel on the right shows the information the user gets when clicking on a gene track. **(b)** The *RHD*/*RHCE* locus as a force-directed layout (top) and a linear layout over CHM13 (bottom), nodes colored by gene, with the edges of an RhD-negative haplotype highlighted (skipping the *RHD* gene loop). **(c)** A subgraph of 949 *E. coli* assemblies de Bruijn graph produced by `Bifrost`. Red nodes are resistance-associated (*f*_*R*_ *>* 0.25 for both drugs; Methods), and colored edges highlight the graph alignments of individual held-out genomes.

`gfaidx` is not limited to pangenome graphs. To show this, we applied it to a bacterial colored de Bruijn graph, built with `Bifrost` (29) from 949 resistant and susceptible *Escherichia coli* assemblies, with nodes colored by their association with antimicrobial resistance (Methods). `gfaidx` indexed this graph in about 3 minutes using only 1 GB of memory. Around a resistance-associated node in the *blaCTX-M-15* region, the extracted subgraph (Fig. 2c) shows that all aligned genomes traversing the region are ampicillin-resistant but only some are ciprofloxacin-resistant. This is consistent with CTX-M-15 being an extended-spectrum *β*-lactamase that hydrolyses ampicillin directly, while its ciprofloxacin association instead reflects co-resistance (30, 31).

In summary, `gfaidx` is an on-disk index for genome graphs whose query cost scales with the region requested rather than with the size of the whole graph. It operates directly on the GFA and is agnostic to how the graph was built, so one index serves pangenome, de Bruijn and assembly graphs alike, and even human pangenome-scale graphs can be indexed and queried on a personal computer. Moreover, the index size is similar to that of the gzip-compressed GFA. For pangenome graphs, it calculates and returns the full set of haplotypes traversing a query with all their sub-coordinates, making it easy to investigate important loci and variations on the graph. Because queries are fast and computationally inexpensive, a single indexed graph can serve many users at once. We therefore host all indexed HPRC graphs behind an API and expose them through `gfaidx-viz,` a WebAssembly visualizer in which users can explore any region of these graphs interactively, with no local download or indexing. `gfaidx’s` libraries further let it be embedded in existing software, as we demon-strate this by integrating the `gfaidx` library in `BandageNG` and Python package in `BubbleGun,` allowing both to operate independently from the graph size, and dependent only on the region they work on at a time. By decoupling subgraph query from the size of the graph, `gfaidx` makes pangenome-scale genome graphs practical practical to explore as they continue to grow.

## Methods

### The gfaidx index

`gfaidx` can index any genome graph type in the GFA 1.x format, treating the GFA records themselves as the object to be indexed, without converting them to a particular variationgraph model, or requiring paths or walks to be present.

#### Partitioning and reordering

`index_gfa` first builds a graph from the node edges defined by the `L` records and applies the Louvain method to partition the graph into communities. Louvain greedily maximizes modularity, grouping densely connected nodes together while minimizing the number of edges that cross between communities. The resulting partition can then be adjusted according to a user-defined size range set by two parameters, a maximum and a minimum number of nodes per community (`–max_chunk_nodes and –min_chunk_nodes`). Communities larger than the maximum are refined by re-running Louvain on their induced subgraph, splitting them into smaller communities. Communities smaller than the minimum are then merged into a neighboring community. However, a merge is applied only when the combined community would still not exceed the maximum, and if no such neighbor exists the small community is left unchanged. Consequently, merging can never create an over-sized community. When several candidate neighbors are available, a community is merged with the one it shares the most edges with, with ties broken toward the smaller resulting community and then by community identifier. This is only run once after the Louvain method, as we found that for the graphs we tested (pangenomes and DBG), one refinement run was more than enough to get balanced community sizes. If no maximum or minimum is given, then `gfaidx` runs the Louvain method iteratively until modularity is maximized. The reason we added the option for an upper and a lower bound for a community, is that an upper bound keeps a small query from having to decompress one very large community, while a lower bound keeps a query from having to open and decompress many tiny communities.

The GFA is then rewritten so that all records of a community are contiguous in the file and compressed as one chunk; the new GFA becomes a multi-member gzip archive, with the shared edges between communities in their own chunk. Because gzip members are independently decompressible, any chunk can be extracted in isolation once its byte range is known.

#### Node identifiers

Each node is assigned a dense integer identifier 0, …, *n* 1, which all indexes below use. Nodes are sorted by community, so that every community owns a con-tiguous range of identifiers, and within a community by a locality key, so that nodes that follow each other along paths receive close identifiers. The locality key is the segment name read as an integer when all names are integers or share a non-numeric prefix followed by digits (e.g., minigraph’s `s123`), since graph builders such as PGGB and Minigraph-Cactus number segments along the paths; otherwise, it is the order in which nodes first appear in the L records.

#### Graph indexing

Alongside the reordered graph`(.gz),index_gfa` generates two sidecars to gain random access through the reordered file: a mapping between each community and its gzip byte offsets `(.idx)`, and a binary hash table mapping every node string identifier to its integer identifier (.ndx). The. `ndx` is a table of 16-byte entries sorted by a 64-bit and a 32-bit FNV-1a hash of the segment name, followed by the first identifier of every community. Given a starting node, `gfaidx` hashes the string ID and binary-searches the table on disk to get the node’s identifier, and then its community by a second binary search over the community start identifiers. Using the community ID, `gfaidx` can get the gzip member byte offsets to decompress that community quickly and parse it.

#### Indexing paths and walks

Paths and walks are stored in a separate index`(.pdx)` that provides two complementary views of the same data, both accessible directly from disk with byte ranges. In the *path-first* view, a path table holds one record per path or walk (its type, name, and the position `step_begin` and length `step_count` of its steps), and a step table stores the steps of all paths back to back. In the *node-first* view, each node record points to its postings, the (path, step) pairs recording which paths visit the node and at which steps. The path-first view enables retrieval of a path’s traversal through the graph, while the node-first view enables retrieval of which path(s) cross a given node and at which step(s), both without materializing the graph in memory.

Both views are compressed by exploiting the locality of the node identifiers. Each step is stored as the difference from the previous step’s identifier together with its orientation,LEB128(2 ·zigzag(*id*_*i*_ −*id*_*i−*1_) + *rev*_*i*_), so a step that moves at most 31 identifiers forwards or 32 backwards takes one byte.

The step table is divided into blocks of 128 steps, each starting from an absolute identifier, and a block index stores the byte offset of every block, so any step is found by reading its block and decoding it up to that step. Node postings are encoded in blocks of 32 consecutive nodes, in increasing identifier order. While encoding a block, `gfaidx` keeps a set of *slots*: for every path that has visited a node of the block so far, the slots hold all of that path’s occurrences (*p, r*) at the most recent such node. An occurrence (*p, r*) at the current node is *predicted* when (*p, r* −1) is a slot, i.e., path *p* arrived directly from that node. Each node is written as the subset of slots it predicts,using the smallest of five descriptions (*none, all*, a bitmap, a list of slot indices, or a list of the slots not predicted), followed by the remaining *explicit* occurrences, stored as gaps in path and step, with the step relative to the path’s last slot when it has one. After a node is written, the paths visiting it replace their slots with their occurrences there, while paths that skip it keep theirs. A predicted occurrence therefore costs at most one bit, a path that takes the other branch of a bubble is predicted again where the branches rejoin, and, because every occurrence has its own slot, a path visiting a region several times, e.g., a copy-number repeat, is predicted on every copy. The first node of each block starts with no slots, and every block can be decoded on its own, so reading a step or the postings of a node takes two small reads and decoding at most one block.

To turn steps into genomic coordinates, a node-length table`(.lnx)` stores one length per node, and a checkpoint file `(.pcx)` stores cumulative path lengths (bases preceding a step) at a regular step interval *N* with a default value of 4,096, and can be changed by the user. The base coordinate of step *b* on a path is then obtained in near-constant time by reading the checkpoint ⌊*b/N*⌋ and summing the lengths of at most *N* −1 remaining steps from. `lnx`. If. `pcx` is absent or fails valida-tion, `gfaidx` falls back to summing the path prefix from the first step on the fly. Optionally, `gfaidx` can build a reference-coordinate index `(.cdx)` separately with `index_coordinates` for a chosen set of paths (usually the reference path). This index maps genomic base-pair intervals along that chosen reference to the graph nodes and reference steps that span them. Because it is a sorted table, a genomic region query `(chr:start–end)` is resolved by binary search directly to the spanning nodes, without walking the reference path, making queries over the indexed reference path(s) fast.

#### Queries

`gfaidx` supports several modes of querying, and for each query mode, it reduces the problem to resolving the targets of the query — whether it was a subgraph, a set of nodes or a subpath — to a set of nodes. `gfaidx` then resolves the nodes to their identifiers through `.ndx,` maps those identifiers to communities, looks up the communities’ byte ranges in `.idx,` and decompresses only those compressed members. `get_subgraph` finds a neighborhood using breadth-first search from one or more starting nodes, then, using the node-first postings in the. `pdx,` it obtains every path and walk crossing the neighborhood nodes. `get_region` resolves a genomic interval through the optional reference-coordinate index .cdx to the set of nodes (anchors) spanning this interval, uses the postings to identify every haplotype crossing them, and for non-reference paths extends each walk between its outermost anchored steps. Coordinates are computed per walk and the required communities are decompressed to emit the matching `S` and `L` records. Output can be restricted to the reference path or expanded to all haplotypes. Alternatively, the region can be defined by distance with the `context` parameter, following `GBZ-base` method: a shortest-path search from the nodes on the reference interval adds every node within `context` base pairs.

#### Access cost

Compression replaces direct addressing of steps and postings with a bounded amount of decoding. Locating step *s* reads one block-index record and the block containing *s*, and decodes at most its 128 values (at most 640 bytes, usually far less than one 4 KiB page). Reading *c* consecutive steps, such as a subpath, decodes at most *c* + 127 values, so a path of *L* steps is read in *O*(*L*) time. Locating the postings of node *v* reads the node records and the encoded bytes of its 32-node block and decodes the block from its first node up to *v*, i.e., at most the postings of one block. Queries visit their nodes in increasing identifier order, so every touched block is decoded once. In the worst case, each requested node is the last of a different block and up to 32 times as many postings are decoded as requested; subgraph and region queries cover contiguous ranges of identifiers and are close to the best case, in which every decoded posting is needed. The amount read and decoded therefore grows with the nodes, steps and communities a query touches.

#### Interfaces

`gfaidx` is distributed as a command-line tool, a C++ library, and a Python package `(pygfaidx)` exposing an immutable `IndexedGraph` object and a mutable Graph object. An `IndexedGraph` can stay open across many queries, so a long-running process or server answering successive requests can run without reopening or rebuilding any structure. Prebuilt statically linked binaries are provided for x86-64 and ARM64. These interfaces allow users to implement their own software on a `gfaidx`-indexed graph.

### Datasets

The HPRC v2 Minigraph-Cactus chromosome 1 graph and the complete CHM13-based HPRC v2 Minigraph-Cactus graph (233 samples, 464 haplotypes) were obtained from the links listed in https://github.com/human-pangenomics/hpp_pangenome_resources. The *E. coli* genome assemblies and laboratory antimicrobial-resistance phenotypes were obtained from the Bacterial and Viral Bioinformatics Resource Center (BV-BRC) (32).

### Tools and versions

Two performance benchmarks were run. The four-tool comparison on chromosome 1 (Fig. 1c) used `gfaidx` v2.3.0, `vg` v1.73.0 (“Ducky”), ODGI v0.9.4 and `GBZ-base` v0.5.1. The whole-genome interval sweep (Fig. 1d) used `gfaidx` v2.3.0 and `GBZ-base` v0.5.1.

### Query selection

For the chromosome 1 comparison, the interval queries were randomly selected, however, avoiding telomeric regions. For the whole-genome interval benchmark, we queried CHM13 intervals of 1, 10 and 100 kb and 1, 2, 5 and 10 Mb centered on 12 loci (84 queries): three centromeric *α*-satellite higher-order-repeat arrays (chr7, chr17, chr22), the MHC, KIR and IGK loci, five structurally complex genes (*AMY1A, SMN1, LPA, KANSL1, CYP2D6*) and a control region on chr3. Each tool extracted the subgraph with 10 kb of context, either as the reference path only `(GBZ-base –reference-only versus gfaidx –mode reference –with_coords)` or as all haplo-types `(GBZ-base versus gfaidx –mode bfs,` with and without– `with_coords`).

### Commands used

#### Indexing

For indexing chromosome 1 HPRC MC graph, we first converted the W lines to P lines for `ODGI` then ran “ `odgi build -g in_graph.gfa -o out_graph.og -O”,` for vg we ran “ `vg convert -g -x in_graph.gfa –ref-sample CHM13”,` for `gfaidx` we ran “ `gfaidx index_gfa in_graph.gfa out_graph.indexed.gfa.gz –progress_every 0 –checkpoint_steps 512 –max_chunk_nodes 10000 –min_chunk_nodes 1000”,` and to construct the `GBZ-base` database, we first needed to run “ `vg gbwt -g out_graph.gbz -G in_graph.gfa –max-node` 0” then “ `gbz-base construct –output out_graph.gbz.db out_graph.gbz”.`

#### Querying

For querying, the following commands were used, for `ODGI` we used “ `odgi extract -i out_graph.og -o subgraph.og -r ’CHM13#0#chr1:$START-$END’ -t 1`” (and `-t 8`), for `vg` we ran “ `vg chunk -x in_graph.xg -p ’CHM13#0#chr1:$START-$END’ –c 0 -O gfa -t 1”,` for `gfaidx` we ran “ `gfaidx get_region out_graph.indexed.gfa.gz chr1:$START-$END subgraph.gfa –reference CHM13 –mode all –with_coords`”.

#### Indexes size

On the HPRC v2 chromosome-1 Minigraph-Cactus graph (7.1 GB as gzip-compressed GFA), the `gfaidx` index occupies 5.1 GB on disk, compared with 50 GB for vg’s xg and 38.5 GB for `ODGI`’s. `og; GBZ-base` is smaller at 1.2 GB (the GBZ together with its SQLite database), reflecting GBZ’s succinct haplotype encoding. Most of `gfaidx`’s footprint (4.5 GB) is the path index. For the complete Minigraph-Cactus graph (66 GB as gzip-compressed GFA), the `gfaidx` index occupies 69 GB, compared with 433 GB and 420 GB for the published vg and `ODGI` graphs (summed over chromosomes) and 18 GB for `GBZ-base;` for the PGGB chromosome-1 graph (7.4 GB as zstd-compressed GFA) it occupies 11 GB, compared with 67 GB (`vg`), 57 GB (`ODGI`) and 7 GB (`GBZ-base`). For graphs without P/W lines, e.g., de Bruijn graphs, the index reduces to the block-compressed GFA file (similar in size to the original compressed GFA, only the lines are re-ordered), plus the community-offset table (.`idx`) which is a few hundred kilobytes and the node hash-map table (.`ndx`) at 16 bytes per node (a 64-bit and a 32-bit string hash plus the 32-bit node identifier), i.e. ~16 MB per million nodes.

### Integrating into BandageNG

To show how `gfaidx` can be utilized in existing software, we integrated its C++ library into `BandageNG,` an interactive viewer for assembly and pangenome graphs derived from`Bandage`. Like `Bandage, BandageNG` parses the entire GFA file into memory before anything can be displayed or extracted. Therefore, inspecting a small region of a large graph requires loading the whole graph first, and graphs larger than the available memory cannot be opened at all on a personal machine. With the integration of `gfaidx, BandageNG` can detect indexed graphs and instead of parsing the file, it opens the index through `gfaidx`’s `IndexedGraph` class. The user then gets to query the graph directly from `BandageNG` using similar queries to the `gfaidx` CLI, namely, `get_subgraph` to get a neighborhood around a node, and `get_region` to get a subgraph around the provided genomic coordinates. With this addition, `BandageNG` can then natively understand and query the indexed graph, allowing users to explore big graphs with-out needing to load the complete graph into memory, and all of `BandageNG`’s functionality works on the subgraph extracted as is without change.

### Integration with BubbleGun

To demonstrate the Python interface, we ran the superbubble detection algorithm from `BubbleGun` on the chromosome 1 Minigraph-Cactus and PGGB graphs of HPRC v2, both in-dexed with `gfaidx. BubbleGun` starts a bounded traversal from every node in both directions and reads the graph only by looking up nodes by ID, so its code could be used unmodified. The in-memory version parses the whole GFA file with its own reader. The indexed version using `pygfaidx` instead fills `BubbleGun`’s table of nodes on demand. It processes the communities in index order. When a traversal reaches a node that is not in memory, it finds the node’s community with `community_id()` and loads that community without sequences with `get_community()`. Cross-community edges are loaded once at the start with `cross_community_edges()`, and when a community is decompressed and loaded, the cross-community edges are also added to it, so a traversal outside the community knows which next community to load when needed. After each traversal, the least recently used communities are dropped until at most 64 remain. Each bubble is found from both of its ends and reported only from the end with the smaller ID, so no global set of visited nodes or found superbubbles is kept.

Both versions reported identical superbubbles (Supplementary Table S2). With the version using `pygfaidx,` peak memory dropped from 12.1 GiB to 1.09 GiB for MC and to 0.52 GiB for PGGB, and the index opened in under 0.2 s instead of the 35–49 s needed to parse the GFA file. Memory depends mainly on the number of cached communities rather than on the size of the graph. This works because few edges cross communities (0.04% in MC, 0.4% in PGGB), so most traversals stay local, i.e., most superbubbles are small and do not traverse different communities. The tradeoff here is time; the indexed version took 1.45 times as long on MC and 5.6 times as long on PGGB, whose traversals span more communities and caused 20,193 community loads, compared with 5,566 for MC. A larger cache trades memory for speed.

### Web application (gfaidx-viz)

The front end was adapted from a prototype of BandageJS (https://github.com/cmdcolin/BandageJS), a browser implementation of the `Bandage` assembly-graph viewer to which multiple features were added. These include native support for GFA P-line and PanSN (33) `W`-line paths; we also added a feature to generate a reference path from minigraph rGFA (`SN,SO,` and `SR` tags) when none is present; a linear layout that pins a chosen reference path to a horizontal axis while keeping force-directed placement for the branches and variants attached to it, this then allows us to have a coordinates ruler and BED-based gene tracks for the chosen path; a copy-number track that bins how often a chosen haplotype revisits each segment of the reference backbone; and a CSV-driven node coloring and labeling similar to the `BandageNG` annotation files.

Graph layout is computed client-side in WebAssembly, using a C++ force-directed algorithm based on the Open Graph Drawing Framework (OGDF) Fast Multipole Multilevel Method (FMMM) (34), to which we added the linear layout. This is compiled with Emscripten into a. `wasm` module that runs in a web worker, so the interface stays responsive while a large graph is laid out. This keeps the compute-intensive layout of large subgraphs entirely in the browser, while the surrounding React/TypeScript application handles parsing, rendering and interaction.

The most important addition, however, is the integration of `gfaidx` into the `Foldseek` webserver that hosts the pre-indexed HPRC graphs. This allows users to query a region of the hosted graphs and receive the requested subgraph. Internally, the API uses `gfaidx` to perform this on the fly. Because only the requested subgraph is ever transferred and drawn, large pre-indexed graphs can be explored interactively by many users without loading the complete graph in the browser.

### Projecting gene annotations per-haplotype onto the graph

Coloring graph nodes by gene normally requires aligning a gene sequence back to the graph, which fails by construction at paralogous loci such as *RHD*/*RHCE* or *C4A*/*C4B*, which are similar enough that an aligner places either sequence on the other’s nodes, and the multi-mappings cannot be differentiated by alignment score only. Previous analyses have handled this by retaining all alignments above a score fraction and inspecting the result manually (35). Here, we avoid alignment entirely. Each assembled haplotype in the HPRC graph is embedded as a `W` walk with information about its sample, haplotype, contig, SeqStart, SeqEnd, and an ordered step list, so accumulating node lengths from `SeqStart` assigns every step a half-open interval in that haplotype’s own contig coordinates. The HPRC per-haplotype gene annotations (CAT GFF3 (36), 464 haplotypes) use the same coordinates, and the annotation seqid plus the PanSN walk name join as `sample#haplotype#seqid;` a gene is then located by binary search over the offset array, returning the step range it occupies on that haplotype’s walk. The only claim made per haplotype is “bases *x*–*y* of this contig are gene *G*” as per the CAT annotation.

### Bacterial de Bruijn graph case study

#### Assembly selection

We downloaded 1,000 *E. coli* assemblies in total from the Bacterial and Viral Bioinformatics Resource Center (32); 500 resistant and 500 susceptible phenotypes to ciprofloxacin and ampicillin that passed assembly quality control (contig count *<* 300 and assembly length between 4.3 and 6.2 Mb). After removing one unavailable assembly (empty FASTA file from the SFTP server) and holding out 50 genomes (25 resistant and 25 susceptible) for alignment, the 949 genomes left were used to build the colored de Bruijn graph.

#### Graph construction

A colored compacted de Bruijn graph (*k* = 31) was built from the 949 assemblies with `Bifrost v1.3.5 (Bifrost build -k 31 -c -r),` which retains every *k*-mer and records the set of genomes containing it. Construction took about 7.5 min and 2 GB of memory and produced 3,528,272 nodes. We then ran `Bifrost query -e 1.0` to get a node/genome presence/absence matrix.

#### Node-phenotype scoring

For a phenotype *p*, let *R*_*p*_ and *S*_*p*_ be the sets of resistant and susceptible genomes used to build the colored de Bruijn graph. For a node *v* and a genome *g*, let

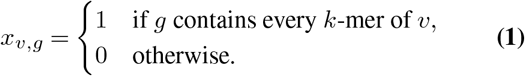

The number of resistant and susceptible genomes containing *v*, and the corresponding fractions, are

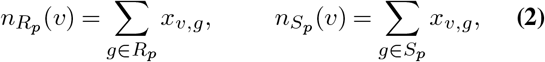

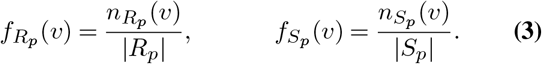

Using the *f*_*R*_(*v*) and *f*_*S*_(*v*) scores, we can color the nodes based on how strongly they are associated with phenotype *p*. The node used as the start for the subgraph extraction in Fig. 2c, within the *blaCTX-M-15* region, had *f*_*R*_ = 0.28 and *f*_*S*_ = 0 for ampicillin and *f*_*R*_ = 0.38 and *f*_*S*_ = 0.016 for ciprofloxacin.

### Aligning the held-out set

We used `GraphAligner` v1.0.20 with the parameter “ `-x dbg`” to align the held-out genomes after cutting them into overlapping 20 kb segments to be interpreted as long reads. We then turned these alignments into W-lines for the visualizer to be able to color the edges as shown in Fig. 2c.

### Hardware

The complete HPRC v2 MC graph benchmark, and `gfaidx` and `GBZ-base` in the chromosome 1 benchmark, were run on a Gigabyte AI TOP ATOM (NVIDIA GB10: a 20-core Arm CPU with 10 Cortex-X925 and 10 Cortex-A725 cores, 128 GB RAM and a 4 TB NVMe SSD). For vg and `ODGI` in the chromosome 1 benchmark, we used an AMD EPYC 7702P 64-Core Processor with 1 TB RAM and 14 TB NVMe scratch storage. For the `BandageNG` chromosome 22 test in Supplementary Figure S2, we used an Apple M4 (4 performance + 6 efficiency cores), 24 GB RAM, macOS personal computer.

## Code availability

gfaidx and gfaidx-viz are free, open-source software. Code can be found at https://github.com/fawaz-dabbaghieh/gfaidx and https://github.com/fawaz-dabbaghieh/graphviz_wasm.

For the BandageNG integration, the source code is at https://github.com/fawaz-dabbaghieh/BandageNG, and for the BubbleGun integration the code is at https://github.com/fawaz-dabbaghieh/gfaidx/tree/main/python/examples/bubblegun.

1. Olson, R. D. *et al*. Introducing the bacterial and viral bioinformatics resource center (BV-BRC): a resource combining PATRIC, IRD and ViPR. *Nucleic Acids Research* **51**, D678–D689 (2023).
2. PanSN-spec contributors. PanSN: Pangenome sequence naming specification. https://github.com/pangenome/PanSN-spec (2024). Accessed 2026.
3. Hachul, S. & Jünger, M. *Drawing Large Graphs with a Potential-Field-Based Multilevel Algorithm*, 285–295 (Springer Berlin Heidelberg, 2005). URL http://dx.doi.org/10.1007/978-3-540-31843-9_29.
4. Liao, W.-W., Asri, M., Ebler, J., Doerr, D. *et al*. A draft human pangenome reference. *Nature* **617**, 312–324 (2023).
5. Fiddes, I. T. *et al*. Comparative Annotation Toolkit (CAT)—simultaneous clade and personal genome annotation. *Genome Research* **28**, 1029–1038 (2018).

## Acknowledgements

We thank C. Marchet for insightful discussions and help in finding biological examples; K. Bonnet for many discussions about the project and the algorithms; P. Heringer on testing `gfaidx` and important user-perspective input; E. Levy Karin (ELKMO) for help revising the manuscript. We acknowledge the support of the Novo Nordisk Foundation (NNF24SA0092560), and InnoCORE program of the Ministry of Science and ICT (No. 1.260017.01).

## Author contributions

**Conceptualization**: F.D. **Data curation**: F.D. **Formal analysis**: F.D. **Methodology**: F.D., M.S. **Resources**: M.S. **Software**: F.D. **Supervision**: M.S. **Writing – original draft**: F.D. **Writing – review & editing**: F.D., M.S.

## Competing interests

M.S. acknowledges outside interest in Stylus Medicine. The remaining authors declare no competing interests.

## References

1. Compeau, P. E. C., Pevzner, P. A. & Tesler, G. How to apply de Bruijn graphs to genome assembly. Nature Biotechnology 29, 987–991 (2011).

2. Cheng, H., Concepcion, G. T., Feng, X., Zhang, H. & Li, H. Haplotype-resolved de novo assembly using phased assembly graphs with hifiasm. Nature Methods 18, 170–175 (2021).

3. Paten, B., Novak, A. M., Eizenga, J. M. & Garrison, E. Genome graphs and the evolution of genome inference. Genome Research 27, 665–676 (2017).

4. Eizenga, J. M., Novak, A. M., Sibbesen, J. A. et al. Pangenome graphs. Annual Review of Genomics and Human Genetics 21, 139–162 (2020).

5. Liao, W.-W., Asri, M., Ebler, J., Doerr, D. et al. A draft human pangenome reference. Nature 617, 312–324 (2023).

6. Lucas, J. K., Hebbar, P., Liao, W.-W., others & Paten, B. Hprc2: A human pangenome reference with near-complete coverage of common genetic variation. bioRxiv (2026). Preprint.

7. Karasikov, M., others & Kahles, A. Efficient and accurate search in petabase-scale sequence repositories. Nature (2025).

8. Jaillard, M. et al. A fast and agnostic method for bacterial genome-wide association studies: Bridging the gap between k-mers and genetic events. PLoS Genetics 14, e1007758 (2018).

9. Li, H. et al. The sequence alignment/map format and samtools. Bioinformatics 25, 2078–2079 (2009).

10. Li, H. Tabix: fast retrieval of sequence features from generic TAB-delimited files. Bioinformatics 27, 718–719 (2011).

11. Danecek, P. et al. Twelve years of SAMtools and BCFtools. GigaScience 10, giab008 (2021).

12. Robinson, J. T. et al. Integrative genomics viewer. Nature Biotechnology 29, 24–26 (2011).

13. Garrison, E. et al. Variation graph toolkit improves read mapping by representing genetic variation in the reference. Nature Biotechnology 36, 875–879 (2018).

14. Hickey, G. et al. Genotyping structural variants in pangenome graphs using the vg toolkit. Genome Biology 21, 35 (2020).

15. Guarracino, A., Heumos, S., Nahnsen, S., Prins, P. & Garrison, E. ODGI: understanding pangenome graphs. Bioinformatics 38, 3319–3326 (2022).

16. Sirén, J. et al. Gbz-base and gaf-base: Indexed pangenome file formats. Bioinformatics (2026).

17. Sirén, J. & Paten, B. GBZ file format for pangenome graphs. Bioinformatics 38, 5012–5018 (2022).

18. GFA-spec Working Group. Graphical fragment assembly (GFA) format specification. https://github.com/GFA-spec/GFA-spec (2024). Accessed 2026.

19. Blondel, V. D., Guillaume, J.-L., Lambiotte, R. & Lefebvre, E. Fast unfolding of communities in large networks. Journal of Statistical Mechanics: Theory and Experiment 2008, P10008 (2008).

20. Wick, R. et al. asl/bandageng. https://github.com/asl/BandageNG.

21. Wick, R. R., Schultz, M. B., Zobel, J. & Holt, K. E. Bandage: interactive visualization of de novo genome assemblies. Bioinformatics 31, 3350–3352 (2015).

22. Dabbaghie, F., Ebler, J. & Marschall, T. BubbleGun: enumerating bubbles and superbubbles in genome graphs. Bioinformatics 38, 4217–4219 (2022).

23. Mirdita, M., Steinegger, M. & Söding, J. MMseqs2 desktop and local web server app for fast, interactive sequence searches. Bioinformatics 35, 2856–2858 (2019).

24. Perry, G. H. et al. Diet and the evolution of human amylase gene copy number variation. Nat. Genet. 39, 1256–1260 (2007).

25. Bolognini, D. et al. Recurrent evolution and selection shape structural diversity at the amylase locus. Nature 634 (2024).

26. Wagner, F. F. & Flegel, W. A. RHD gene deletion occurred in the Rhesus box. Blood 95, 3662–3668 (2000).

27. Yang, Y. et al. Gene copy-number variation and associated polymorphisms of complement component C4 in human systemic lupus erythematosus. American Journal of Human Genetics 80, 1037–1054 (2007).

28. Sekar, A. et al. Schizophrenia risk from complex variation of complement component 4. Nature 530, 177–183 (2016).

29. Holley, G. & Melsted, P. Bifrost: highly parallel construction and indexing of colored and compacted de Bruijn graphs. Genome Biology 21, 249 (2020).

30. Price, L. B., Johnson, J. R., Aziz, M. et al. The epidemic of extended-spectrum-β-lactamase-producing Escherichia coli ST131 is driven by a single highly pathogenic subclone, H30-rx. mBio 4, e00377–13 (2013).

31. Coque, T. M., Novais, Â., Carattoli, A., Poirel, L. et al. Dissemination of clonally related Escherichia coli strains expressing extended-spectrum β-lactamase CTX-M-15. Emerging Infectious Diseases 14, 195–200 (2008).

## References

32. Olson, R. D. et al. Introducing the bacterial and viral bioinformatics resource center (BV-BRC): a resource combining PATRIC, IRD and ViPR. Nucleic Acids Research 51, D678–D689 (2023).

33. PanSN-spec contributors. PanSN: Pangenome sequence naming specification. https://github.com/pangenome/PanSN-spec (2024). Accessed 2026.

34. Hachul, S. & Jünger, M. Drawing Large Graphs with a Potential-Field-Based Multilevel Algorithm, 285–295 (Springer Berlin Heidelberg, 2005). URL 10.1007/978-3-540-31843-9_29.

35. Liao, W.-W., Asri, M., Ebler, J., Doerr, D. et al. A draft human pangenome reference. Nature 617, 312–324 (2023).

36. Fiddes, I. T. et al. Comparative Annotation Toolkit (CAT)—simultaneous clade and personal genome annotation. Genome Research 28, 1029–1038 (2018).

