## supplementary figures for "Efficient genome graphs indexing, querying and visualization with gfaidx and gfaidx-viz"

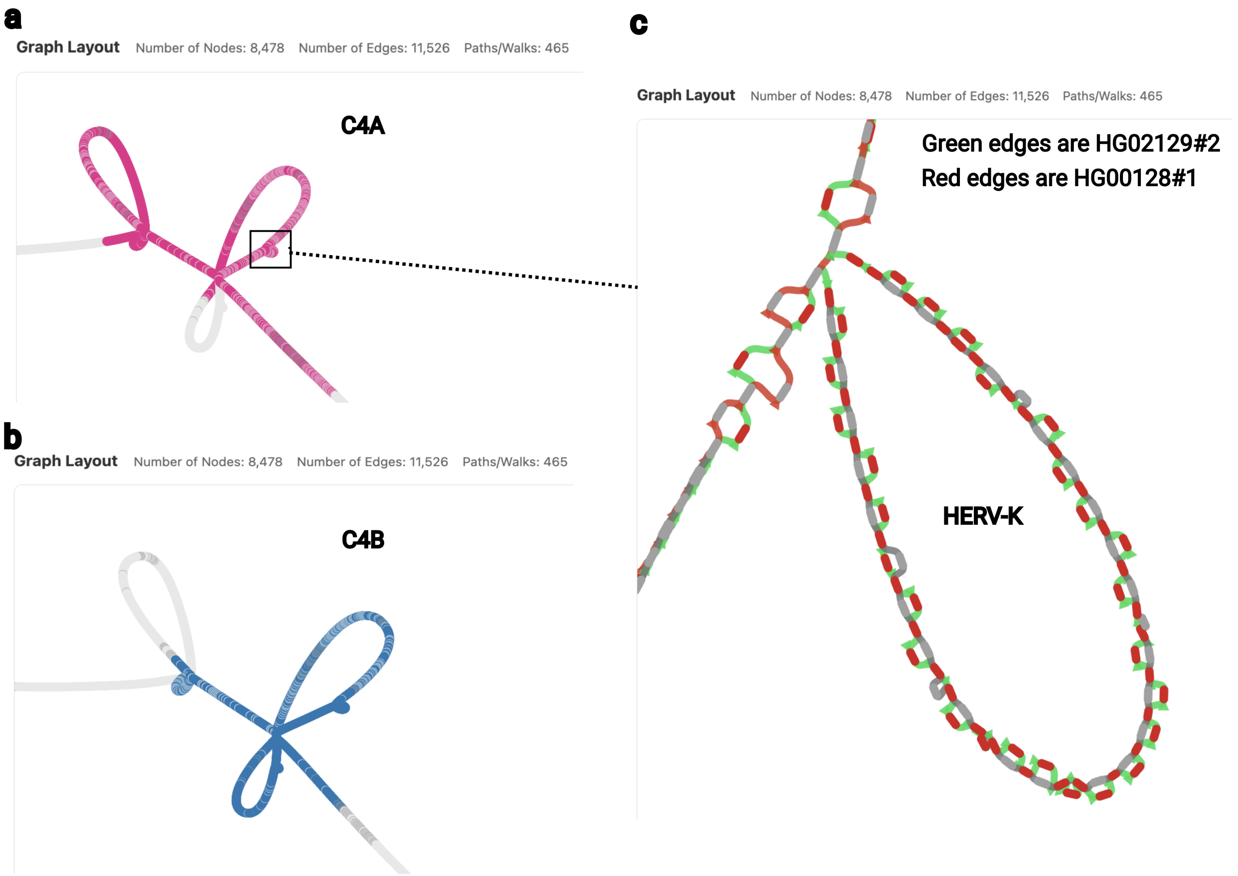


Supplementary Figure S1: **The *C4* (RCCX) locus and the HERV-K(C4) insertion in the HPRC v2 Minigraph-Cactus graph**, produced with `gfaidx` and visualized with `gfaidx-viz`. **(a)** Nodes colored for *C4A* and **(b)** for *C4B*; nodes carrying both colors are those the graph does not resolve between the two paralogs (44% of *C4*-labeled nodes). **(c)** Zoom on the $\sim$6.4 kb HERV-K(C4) insertion in intron 9 (red nodes): the walk of HG00128 haplotype 1 (red edges) takes the short path and bypasses the insertion, whereas HG02129 haplotype 2 (green edges) traverses the HERV-K loop, distinguishing short- and long-*C4* alleles.

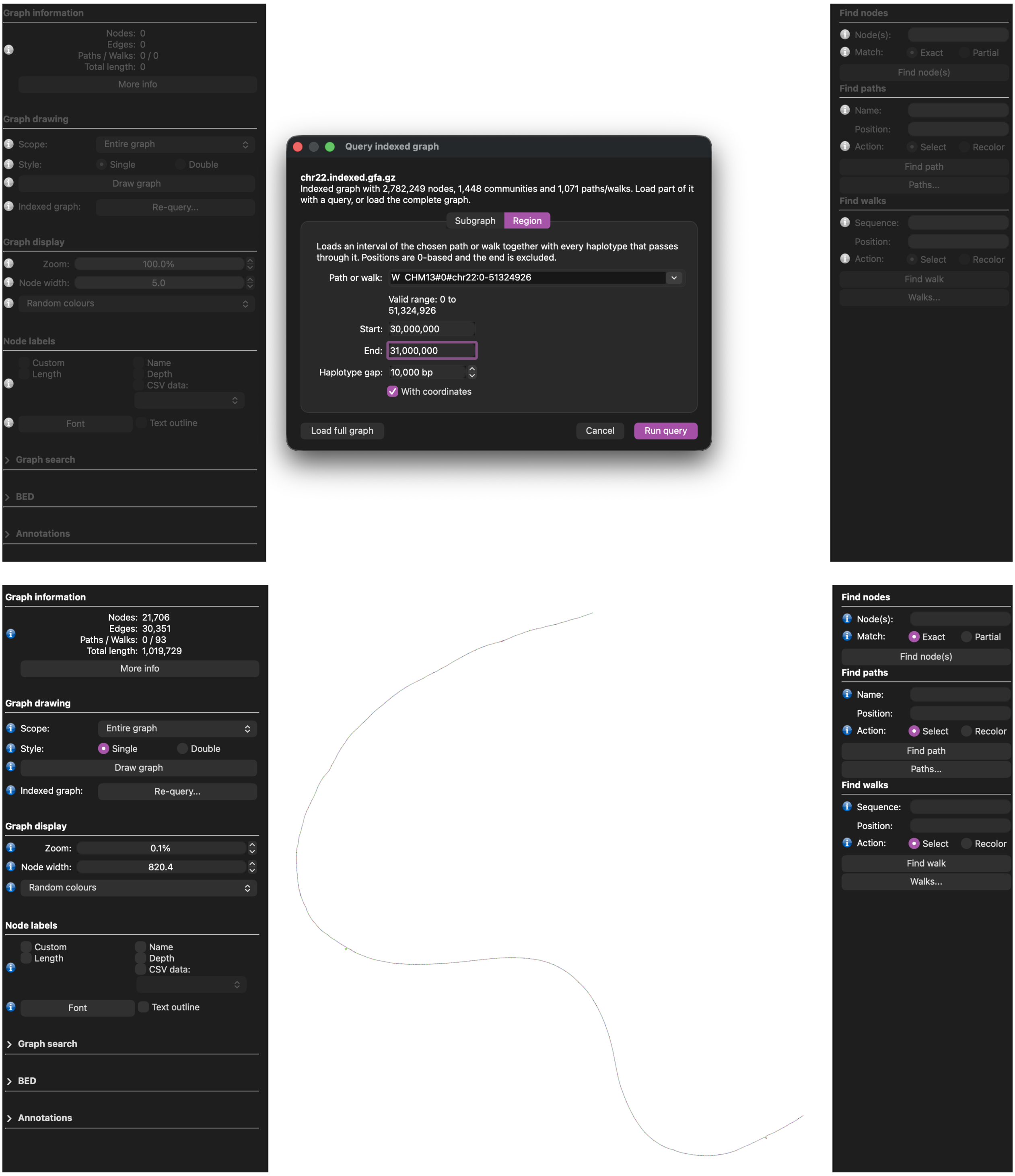


Supplementary Figure S2: **BandageNG with the gfaidx library integration.** We have added a popup window inside the app that shows up when an indexed graph is selected by `BandageNG`. The users then can query a neighborhood around a node, or query genomic intervals to get a subgraph, which then internally gets extracted and loaded into the app for visualization. Users can then click the "Re-query" button to make a new query. This integration not only improves the speed and memory use of `BandageNG`, but also allows it to access and visualize subgraphs of graphs that would otherwise be too big to be loaded in RAM on a personal machine. We tested this with the HPRC v1 Minigraph-Cactus graph of chromosome 22, where we chose a random node and extracted the neighborhood around it using both `BandageNG`'s internal extraction method and `gfaidx`. With the `BandageNG` method, it required 12 seconds and 3 GB of memory to load the complete graph, while with `gfaidx`, it ran the extraction in 0.2 seconds and used only 50 MB.

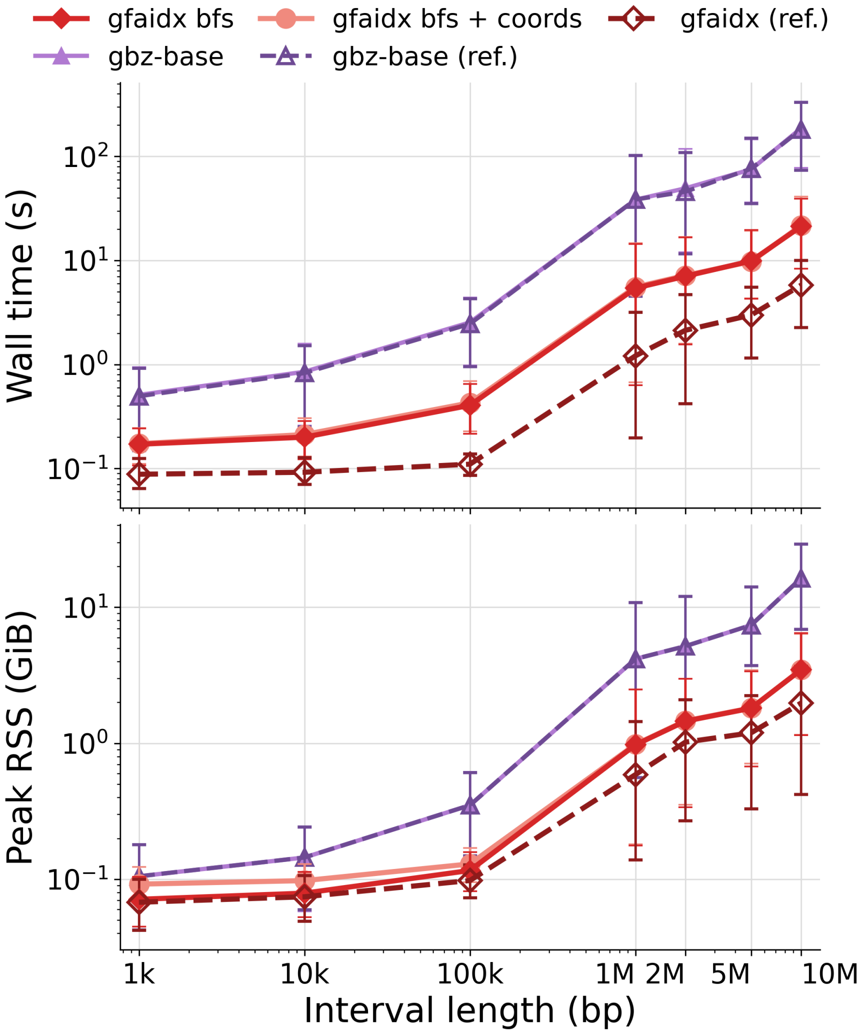


Supplementary Figure S3: **Difficult loci query benchmark on chromosome 1 of the PGGB graph.** Here we compare only `GBZ-base` and `gfaidx` on difficult loci, which were centered on ten CHM13 chr1 loci: the active centromeric HOR array, the 1q12 HSat2 array, the 1q21.1 *NOTCH2NL/NBPF* segmental duplications, *AMY1*, *RHD–RHCE*, *GBA–GBAP1*, *FCGR2A–FCGR2B* and *CFH–CFHR*, plus two random control positions, each at seven interval sizes from 1 kb to 10 Mb. `gfaidx` returned the same node sets as `GBZ-base` for all queries. Its advantage grows with interval size: at 10 Mb it was on average 7× faster for all-haplotype extraction (26 s vs 187 s) and 29× faster for reference-only extraction (6.3 s vs 182 s), with 4–8× lower peak memory (3.4 vs 16.4 GiB and 2.0 vs 16.4 GiB). At 1 kb both tools finish in under a second, and `GBZ-base` is faster for satellite queries that resolve to a single large node.

| | gfaidx | vg | ODGI | GBZ-base |
|---|---|---|---|---|
| Primary focus | Random access / extraction from GFA | Variation-graph toolkit (mapping, genotyping) | Pangenome graph analysis & manipulation | Disk-backed queries on GBZ |
| Native input | GFA 1.0/1.1, rGFA | GFA, GBZ, XG, PackedGraph | .og; GFA 1.0 | GBZ |
| Access model | External memory (disk, block-compressed) | In memory | In memory (succinct) | External memory (SQLite) |
| GFA 1.1 W walks | ✓ | ✓ | ✗ (convert to P) | ✓ (via GBZ) |
| rGFA coordinates | ✓ | partial | ✗ | ✗ |
| Extraction returns all haplotypes | ✓ (named + coords) | ✓ | ✓ | ✓ (unnamed walks) |
| API | C++ & Python | C++ (CLI) | C++ & Python | Rust |

Supplementary Table S1: **Comparing the different features of the tools used:** Capability and focus of `gfaidx` versus three widely used graph toolkits. ✓, supported; ✗, not supported or not applicable.

| | MC | | PGGB | |
|---|---|---|---|---|
| | In memory | pygfaidx | In memory | pygfaidx |
| Load graph / open index (s) | 35.0 | 0.02 | 49.3 | 0.18 |
| Bubble detection (s) | 56.6 | 144.3 | 63.1 | 683.6 |
| Wall time (s) | 99.6 | 144.4 | 122.8 | 683.9 |
| Peak memory (GiB) | 12.11 | 1.09 | 12.15 | 0.52 |
| Bubbles | 3,680,188 | | 3,558,275 | |
| simple | 3,066,604 | | 3,030,501 | |
| insertions | 306,972 | | 264,877 | |
| superbubbles | 306,612 | | 262,897 | |

Supplementary Table S2: Bubble detection on chromosome 1 of the HPRC v2 Minigraph-Cactus (MC) and PGGB graphs with BubbleGun's superbubble algorithm. The "In memory" column shows the results for reading the whole graph into memory, i.e., the default `BubbleGun` behavior. The `pygfaidx` column has the results for reading the indexed graph community by community through the `pygfaidx` Python interface, keeping up to 64 communities cached between searches. Both modes run the same, unmodified bubble detection algorithm, and report identical bubbles. Times are wall-clock seconds and memory is the peak resident set size, both from `gnu time -v`.
